# The transmembrane protein TMEM127 regulates activation and fate of an MHC-I degradation complex by WWP2 modifications

**DOI:** 10.64898/2026.06.29.735340

**Authors:** Hector Gonzalez-Cantu, Viviane Nascimento da Conceicao, Syeda Yusra Munawar, Karen Johns, Carine Jaafar, Nadia Reyna, Arshjot Multani, Cynthia M. Estrada-Zuniga, Daohong Zhou, Ricardo C. T. Aguiar, Yaxia Yuan, Patricia L. M. Dahia

**Affiliations:** Division of Hematology-Medical Oncology, Department of Medicine, University of Texas Health San Antonio (UTHSA); Department of Biochemistry & Structural Biology; Mays Cancer Center at UTHSA; South Texas Veterans Health Care System, Audie Murphy Memorial Veterans Hospital, San Antonio, TX)

**Keywords:** TMEM127, MHC-I, WWP2, HECT E3 ligase, ubiquitination, endocytosis, PxxY motif, antigen presentation, adaptive immunity, degradation

## Abstract

TMEM127 is an adaptor protein that bridges substrates to E3 ubiquitin ligases of the HECT family. Among its interacting partners is the major histocompatibility class I (MHC-I), a critical component of the antigen presentation pathway and the adaptive immune response. MHC-I is ubiquitinated and fated for lysosome-mediated degradation by the WWP2 E3 ligase in a complex that involves TMEM127 and a second adaptor protein, SUSD6. However, the interacting dynamics among complex components remains to be determined, a key knowledge gap towards the development of pharmacological modulators. Here, using *in vitro* and *in vivo* models, we report that TMEM127-WWP2 interaction stabilizes the MHC-I degradation complex and reveals an asymmetric role of the two adaptor proteins. Specifically, we find that TMEM127 regulates WWP2 catalytic activity, abundance and localization through its canonical PY motif interaction with the WW domain of WWP2 with contribution of a TMEM127 endocytic motif, providing a mechanism to restrain complex activity. Further, we validate the impact of TMEM127 dosage in the endogenous complex assembly and regulation. Our results nominate TMEM127 as a critical member of the MHC-I degradation complex and highlight the TMEM127-WWP2 interaction as a target for augmenting MHC-I-mediated antigen presentation, a long sought goal in cancer immunology.

## Introduction

We previously identified germline loss-of-function (LOF) mutations of the ubiquitously expressed *TMEM127* gene as causative drivers of pheochromocytomas and paragangliomas (PPGL), rare neuroendocrine tumors of the adrenal medulla or paraganglia(*1*). Modeling tumor-associated *TMEM127* variants revealed disruption of the TMEM127 subcellular distribution, from a predominant endomembrane, punctate pattern in favor of a diffuse, cytosolic distribution, suggesting that the TMEM127 cellular function is dependent on its endomembrane trafficking abilities(*1–5*). In addition, we uncovered a four-transmembrane architecture and the presence of a functional endocytic domain in its C-terminus (*5*). An important insight into TMEM127 function emerged in a distinct context, when it was identified as an obligatory modulator of major histocompatibility class II (MHC-II) in dendritic cells infected with Salmonella through TMEM127’s association with the WWP2 HECT E3 ubiquitin ligase (*6*). Using in vitro and in vivo adrenomedullary cell models, we found that TMEM127 regulates proliferative signals by antagonizing the expression of cell surface RET kinase receptor, which can be oncogenic in adrenomedullary tissue, via recruitment of NEDD4, the prototypical HECT E3 ubiquitin ligase (*7*). We found that TMEM127 binds to RET and bridges it to NEDD4 via a conserved proline-rich domain (PxxY or PY motif) located at TMEM127 C-terminus. The NEDD4-TMEM127 complex promotes RET ubiquitination, endocytosis, and lysosomal degradation(*8*). In agreement with the translational relevance of these findings, deficiency of TMEM127 in adrenomedullary tumors confers high sensitivity to selective RET inhibition in vivo (*8*).

TMEM127 was also implicated, through unbiased screens, in pathways that govern antigen presentation and MHC-I (*9, 10*). MHC-I is a ubiquitously expressed dimeric heterocomplex comprised of a variable human leukocytic antigen (HLA) heavy chain unit and the invariable beta 2 microglobulin (B2M) unit, serving a key role in the antigen presentation pathway of the adaptive immune system. It regulates immune surveillance by presenting cytosolic antigens to circulating CD8+ T lymphocytes, where a ‘normal’ self-antigen signals the healthy status of the cell, while a tumor-associated antigen signals oncogenic transformation, triggering a CD8+-dependent cytotoxic response(*11–13*). Antigen presentation by cancer cells is at the crux of efficient antitumor immune surveillance responses(*11*). Cancer cells can increase their fitness by dampening antigen presentation through mechanisms that include, among others, downregulation of cell surface MHC-I(*14–16*). Improving and retaining MHC-I surface expression and antigen presentation has been recognized as an effective strategy to improve anti-tumor immune responses, with far reaching implications in oncology(*17, 18*). Chen and colleagues discovered that surface anchored MHC-I was targeted by a multi-protein degradation complex composed of the transmembrane proteins **<u>S</u>**USD6 and **<u>T</u>**MEM127, and the HECT E3 ubiquitin ligase **<u>W</u>**WP2 (**STW**) (*10*). The STW complex mediates MHC-I ubiquitination, internalization, and lysosome-mediated degradation. Conversely, tumor cell models deficient for *Susd6* or *Tmem127* increased MHC-I cell surface density and resulted in improved CD8+ T cell-dependent antitumor responses, supporting the biological relevance of this complex to curb MHC-I activity, and establishing STW as a potential target to augment antitumor immune surveillance(*10*).

The close parallels presented by the three TMEM127-related E3 ligase complexes described above suggest that they may share organizational and regulatory attributes. However, this information remains largely undetermined. In particular, understanding how the STW axis functionally regulates MHC-I suppression will inform future studies that aim to target the complex to increase MHC-I expression and antigen presentation in cancer. Here, we investigated the contribution of TMEM127 to the STW complex. Our findings delineate a dominant stabilizing role of TMEM127 toward STW and uncover a regulatory impact of TMEM127 on WWP2-mediated ubiquitination, which may confer a mechanism to self-limit STW activity. We also uncover vulnerabilities that could be potentially targetable for translational purposes.

## Results

### MHC-I recruitment to degradation complex is dependent on TMEM127

To gain insights into the STW complex structure, we generated a 3D rendition of MHC-I in complex with STW using AlphaFold 3 (Fig 1A)(*19*). This model predicted that TMEM127 was positioned centrally within the complex, with its four transmembrane domains flanked by the single transmembrane domains of SUSD6 and MHC-I (i.e., HLA/B2M), respectively, while WWP2 was predicted to interface with cytosolic regions of SUSD6, TMEM127, and MHC-I. Guided by this structural model, we investigated if TMEM127 interacted with WWP2, SUSD6, or MHC-I by co-expressing DNA constructs coding for WWP2-Myc/Flag, HLA-B (a haplotype of *HLA*), Flag-SUSD6, and HA-TMEM127, either individually or in combination, and performing an HA (TMEM127) co-immunoprecipitation assay. For these experiments, we created HEK293 cells genetically doubly depleted of *WWP2* and *TMEM127*, which also have undetectable SUSD6 expression, thus minimizing background expression of endogenous STW proteins (Suppl. Fig 1A). We found that TMEM127 co-immunoprecipitated WWP2, SUSD6, and HLA, in agreement with the prediction (Fig 1B, Suppl Fig 1B). In the original description of the STW model, genetic depletion of *Susd6* or *Tmem127* resulted in increased MHC-I levels, suggesting that these adaptor proteins were necessary for WWP2-MHC-I interactions and that their absence resulted in MHC-I accumulation(*10*). To address this premise, we co-immunoprecipitated WWP2 (Myc tag) in cell lysates from HEK293 cells co-expressing relevant DNA constructs. These assays revealed that WWP2-MHC-I interactions were detectable in protein lysates from cells co-expressing TMEM127 and SUSD6 (Fig1C, lane 2). In addition, WWP2-MHC-I interactions were still detectable in the presence of either TMEM127 or SUSD6 alone (Fig 1C, lanes 3, and 4), but not in cells lacking both TMEM127 and SUSD6 constructs, suggesting that the expression of either adaptor was sufficient to bridge WWP2-MHC-I interactions(Fig 1C, lane 5), as predicted by the 3D model. Reciprocally, MHC-I co-immunoprecipitations confirmed that the interaction between HLA and WWP2 was dependent on TMEM127 or SUSD6, in further agreement with the model (Supp. Fig 1C).

**Figure 1.**
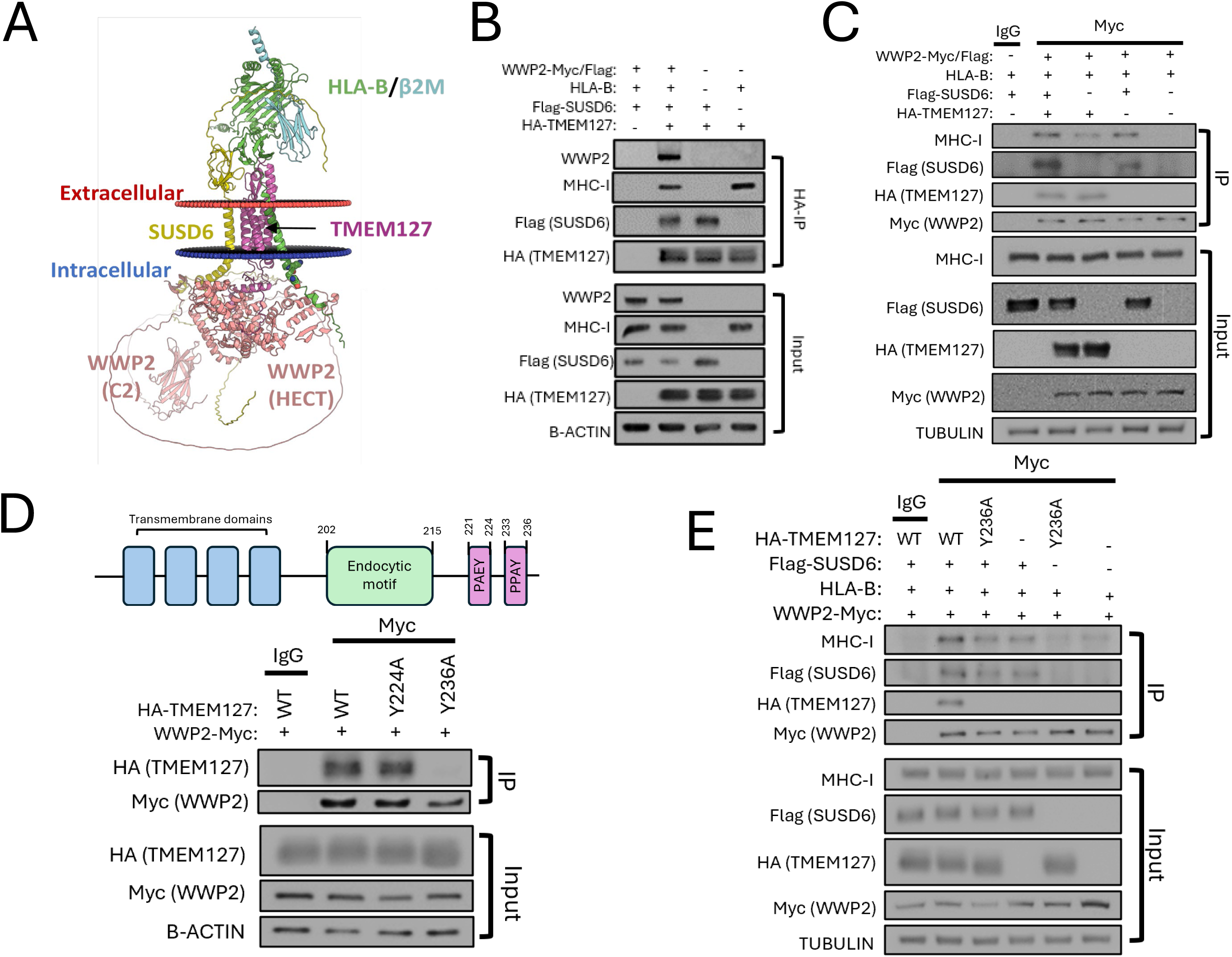
MHC-I recruitment to the STW degradation complex is dependent on TMEM127. **A)** Modeling of TMEM127 by Alpha-Fold3 with SUSD6, WWP2, and MHC-I (including HLA-B and Beta 2 microglobulin (B2M); **B)** Protein lysates from *WWP2*- and *TMEM127*-depleted HEK293 cells co-transfected with WWP2-Myc/Flag, HLA-B, Flag-SUSD6, and HA-TMEM127 were co-immunoprecipitated with an HA (TMEM127) or an IgG serotype control antibody and immunoprecipitates were immunoblotted for WWP2, Flag (SUSD6), and MHC-I. Corresponding whole cell lysate inputs were analyzed by immunoblotting with identical antibodies as co-IP blots. N=3; **C)** Protein lysates from WWP2 and TMEM127-depleted HEK293 cells co-transfected with WWP2-Myc/Flag, HLA-B, Flag-SUSD6, and HA-TMEM127 were co-immunoprecipitated with a Myc (WWP2) or an IgG serotype control antibody and immunoprecipitates were immunoblotted for MHC-I, Flag (SUSD6), HA (TMEM127). Corresponding whole cell lysate inputs were analyzed by immunoblotting with identical antibodies as co-IP blots. N=2; **D)** (Top) Graphical representation of TMEM127 structural and transmembrane domains, endocytic domain, non-canonical PY motif, and canonical PY motif; (Bottom) Protein lysates from WWP2 and TMEM127-depleted HEK293 cells co-transfected with WWP2-Myc and HA-TMEM127 WT, or Y224A and Y236A mutants were co-immunoprecipitated with a Myc (WWP2) or an IgG serotype control antibody and immunoprecipitates were immunoblotted for HA (TMEM127). N=3; **E)** Protein lysates from WWP2 and TMEM127-depleted HEK293 cells co-transfected with WWP2-Myc/Flag, HLA-B, Flag-SUSD6, and HA-TMEM127 WT or Y236A mutant were co-immunoprecipitated with a Myc (WWP2) or an IgG serotype control antibody and immunoprecipitates were immunoblotted for MHC-I, Flag (SUSD6), HA (TMEM127). Corresponding whole cell lysate inputs were analyzed by immunoblotting with identical antibodies as co-IP blots. N=2.

We expanded on these findings by asking whether specifically abolishing TMEM127-WWP2 interactions could impact the STW complex. First, using a TMEM127 (Y236A) mutant, we showed that the physical interaction between TMEM127 and WWP2 is dependent on the TMEM127 C-terminus PY motif (PPAY), analogous to our previously reported observations in the context of TMEM127-NEDD4 interaction and in an independent model (*6, 8*), while an upstream, noncanonical PY-like motif (PAEY), maintained WWP2 binding (Fig 1D). Next, to directly compare the impact of TMEM127 Y236A on the STW complex assembly, we co-immunoprecipitated WWP2 from lysates of HEK293 cells co-expressing various construct combinations. These experiments revealed that the interaction between WWP2 and HLA was negligible in TMEM127 knockout (KO) cells expressing vector only or TMEM127 Y236A only (Fig 1E), but it was detectable, although to a lesser extent than in TMEM127 WT cells, in samples co-expressing SUSD6 in combination with TMEM127 Y236A (Fig 1E). When taken together with our earlier results (Fig 1C), these findings suggested that TMEM127 and/or SUSD6 were necessary to bridge the physical interaction between WWP2 and HLA, and that abolishing TMEM127-WWP2 binding weakened or completely ablated (TMEM127 Y236A expression without SUSD6 expression) the WWP2-HLA interaction.

To connect these findings with the complex function, we next analyzed ubiquitination signals from lysates of HEK293 cells co-expressing WWP2, TMEM127, and HA-ubiquitin using MHC-I pulldown assays. We found that TMEM127 re-expression led to increased HLA ubiquitination relative to empty vector (EV) control samples (Fig 2A). We also examined the impact of SUSD6 by adding it alone or combination with TMEM127. Expression of SUSD6 alone did not elicit ubiquitination signals from HLA, and co-expression of TMEM127 and SUSD6 elicited ubiquitination signals from HLA similar in magnitude to that of TMEM127 alone (Fig 2B), suggesting that HLA ubiquitination by STW was dominantly linked to TMEM127.

**Figure 2.**
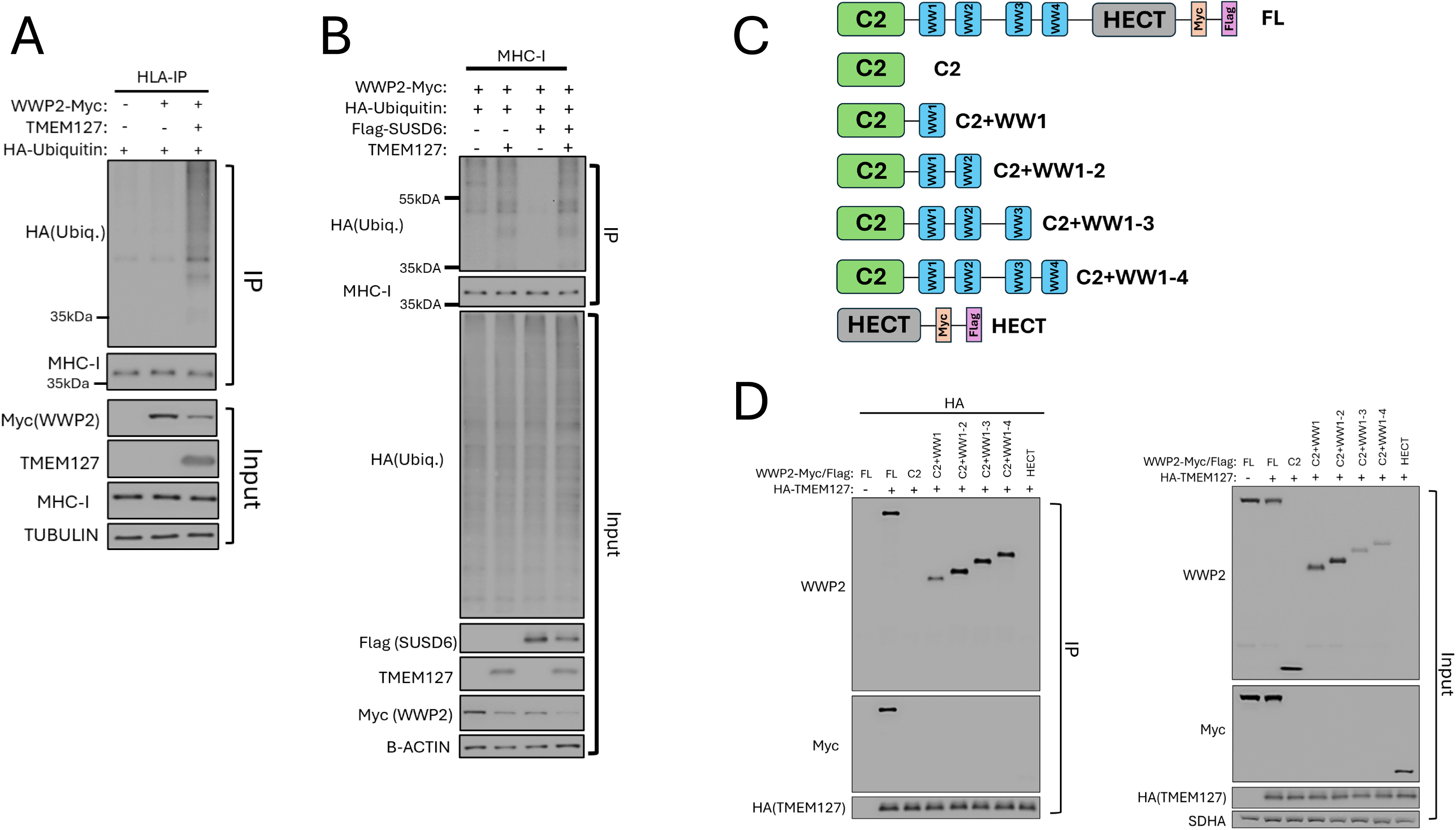
MHC-I ubiquitination is dependent on functional TMEM127-WWP2 interactions. **A)** Protein lysates from WWP2 and TMEM127-depleted HEK293 cells co-transfected with WWP2-Myc/Flag, HLA-B, HA-ubiquitin, and untagged TMEM127 were co-immunoprecipitated with an MHC-I or an IgG serotype control antibody and immunoprecipitates were immunoblotted for MHC-I, and HA (ubiquitin). Corresponding whole cell lysate inputs were analyzed by immunoblotting with identical antibodies as co-IP blots. N=2; **B)** Protein lysates from WWP2 and TMEM127-depleted HEK293 cells co-transfected with WWP2-Myc, HA-ubiquitin, Flag-SUSD6 and untagged TMEM127 were co-immunoprecipitated with an MHC-I antibody and immunoprecipitates were immunoblotted for MHC-I, and HA (ubiquitin). Corresponding whole cell lysate inputs were analyzed by immunoblotting with identical antibodies as co-IP blots. N=4; **C)** Graphical representation of WWP2 truncation constructs. Full-length (FL) and HECT domain (HECT) constructs retained the dual C-terminus Myc/Flag epitope tag. ‘C2’ contains the C2 N-terminus domain, ‘C2+WW1’ contains the C2 N-terminus domain plus WW1, ‘C2+WW1-2’ contains the C2 N-terminus domain plus WW1 and WW2, ‘C2+WW1-3’ contains the C2 N-terminus domain plus WW1, WW2, and WW3, and ‘C2+WW1-4’ contains the C2 N-terminus domain plus WW1, WW2, WW3, and WW4; **D)** (Left) Protein lysates from WWP2 and TMEM127-depleted HEK293 cells co-transfected with WWP2 truncation constructs and HA-TMEM127 were co-immunoprecipitated with an HA (TMEM127) antibody and immunoprecipitates were immunoblotted for WWP2 and Myc (WWP2 FL, and HECT domain). (Right) Corresponding whole cell lysate inputs were analyzed by immunoblotting with identical antibodies as co-IP blots. N=2.

### TMEM127 promotes WWP2 autoubiquitination and abundance through canonical PY and also endocytic interactions

To map the interactions between TMEM127 and WWP2 and gain insights into their relevance to the complex, we generated recombinant constructs expressing subsets of WWP2 domains: ‘C2’ containing its N-terminus C2 domain, which harbors phospholipid membrane binding properties, and combinations of C2 and the four WW domains (Fig 2C). These domains, characteristic of the HECT family of E3 ligases, comprise two conserved, regularly spaced tryptophan residues and serve as the main binding sites for PY-motif containing substrates and/or binding adaptors. We also generated a ‘HECT’ only construct, consisting of the catalytic domain of WWP2(*20*). We performed HA (TMEM127) co-immunoprecipitation (co-IP) assays with lysates from HEK293 cells co-transfected with the various WWP2 truncations or a full-length (FL) version, and HA-TMEM127. Using WWP2 FL as positive control, these assays revealed that TMEM127 also pulled down all WWP2 WW-domain containing constructs, but not the C2-only or HECT-only domains (Fig 2D). These findings confirm that TMEM127-WWP2 interactions consist exclusively of canonical PY-WW domain interactions and were strongest with constructs that shared the second WW domain (WW2), in agreement with a recent report(*21*), although they were also detectable with a WW1-containing construct (Fig 2D).

In parallel, we observed that the expression of the WWP2 construct was downregulated by TMEM127 alone or by TMEM127+SUSD6 co-expression (Fig 2B, input blot). We considered that this decrease in WWP2 protein levels could be due to WWP2 autoubiquitination, an effect that is often observed in HECT E3 ligase-related catalytic activation(*22, 23*). To investigate this hypothesis, we evaluated WWP2 ubiquitination signals in these samples. These assays revealed that TMEM127 expression alone led to increased WWP2 autoubiquitination signals relative to an EV control (Fig 3A). These signals were more evident in the presence of ectopic HLA-B (Suppl. Fig 2). Combined TMEM127 and SUSD6 co-expression produced weaker WWP2 ubiquitination signals than those seen with expression of TMEM127 alone and SUSD6 ectopic expression alone yielded no detectable WWP2 autoubiquitination (Fig 3A). The ubiquitination signals tracked with WWP2 input levels in each of these samples, suggesting that TMEM127 promoted WWP2 autoubiquitination.

**Figure 3.**
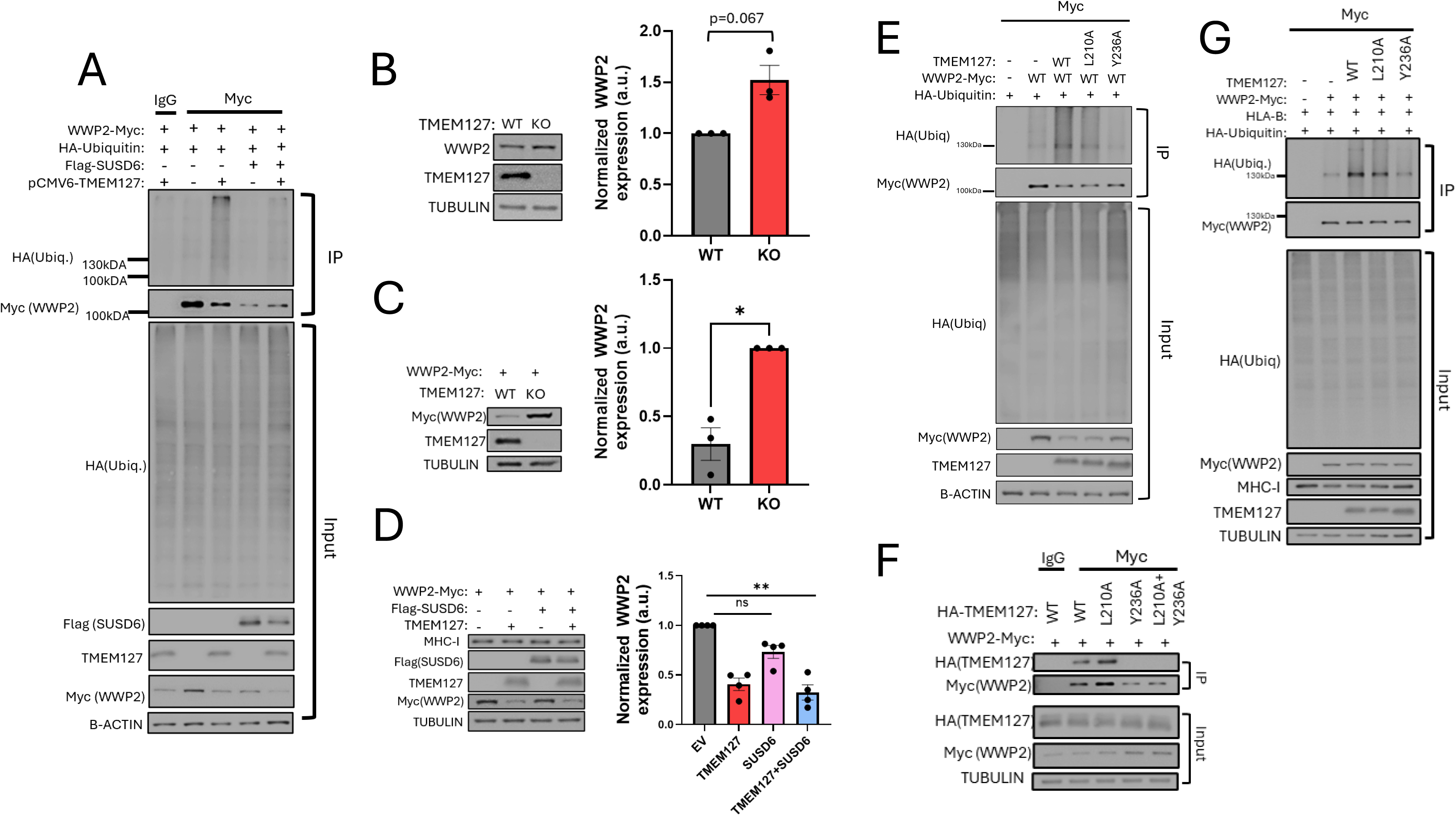
TMEM127 regulates WWP2 autoubiquitination and abundance through PY and endocytic motifs. **A)** Protein lysates from WWP2 and TMEM127-depleted HEK293 cells co-transfected with WWP2-Myc/Flag, HA-ubiquitin, and untagged TMEM127 were co-immunoprecipitated with a Myc (WWP2) or an IgG serotype control antibody and immunoprecipitates were immunoblotted for Myc (WWP2), and HA (ubiquitin). Note, 2C and 2D are reciprocal co-IP experiments. N=4;; **B)** (Left) Protein lysates from TMEM127 KO and TMEM127 WT HEK293 cells were analyzed for endogenous WWP2 expression by immunoblotting with a WWP2 antibody. (Right) Quantification of normalized WWP2 bands by Tubulin loading control (N=3). Data is represented as percent of ‘WT’ and was statistically analyzed using an unpaired t test with Welch’s correction. N=3; **C)** (Left) Protein lysates from double WWP2 KO and TMEM127 KO HEK293 cells (TMEM127 KO) and single WWP2 KO HEK293 cells (TMEM127 WT) ectopically expressing WWP2-Myc were analyzed for WWP2 expression by immunoblotting with a Myc (WWP2) antibody. (Right) Quantification of normalized WWP2 bands by Tubulin loading control (N=3). Data is represented as percent of ‘TMEM127 KO’ and was statistically analyzed using an unpaired t test with Welch’s correction. (*): p<0.05; **D)** (Left) Protein lysates from double WWP2 KO and TMEM127 KO HEK293 cells co-transfected with WWP2-Myc, and untagged TMEM127 or Flag-SUSD6 and immunoblotted for Myc (WWP2), TMEM127, Flag (SUSD6) or Tubulin. (Right) Quantification of normalized WWP2 bands relative to Tubulin (N=4). Data is represented as percent of EV (KO) and was statistically analyzed using ANOVA-Friedman test and Wilcoxon matched-pairs signed rank test (n.s); **E**) Protein lysates from WWP2 and TMEM127-depleted HEK293 cells co-transfected with WWP2-Myc, HA-ubiquitin, and TMEM127 WT and L210A (endocytic mutant) or Y236A (PY mutant) were co-immunoprecipitated with a Myc (WWP2) or an IgG serotype control antibody and immunoprecipitates were immunoblotted for Myc (WWP2), and HA (ubiquitin). Corresponding whole cell lysate inputs were analyzed by immunoblotting with identical antibodies as co-IP blots. N=2; **F)** Protein lysates from WWP2 and TMEM127-depleted HEK293 cells co-transfected with WWP2-Myc and HA-TMEM127 WT, or L210A, Y236A, or L210A/Y236A mutants were co-immunoprecipitated with a Myc (WWP2) or an IgG serotype control antibody and immunoprecipitates were immunoblotted for HA (TMEM127). Corresponding whole cell lysate inputs were analyzed by immunoblotting with identical antibodies as co-IP blots. N=2; **G)** Protein lysates from WWP2 and TMEM127-depleted HEK293 cells co-transfected with WWP2-Myc, HA-ubiquitin, HLA-B, and untagged TMEM127 WT and L210A or Y236A mutants were co-immunoprecipitated with a Myc (WWP2) or an IgG serotype control antibody and immunoprecipitates were immunoblotted for Myc (WWP2), and HA (ubiquitin). Corresponding whole cell lysate inputs were analyzed by immunoblotting with identical antibodies as co-IP blots. N=2.

Next, we sought to gain further insights into TMEM127 impact toward WWP2 abundance. Our HA-ubiquitin co-IP experiments described above suggested that WWP2 expression decreased in a TMEM127-dependent manner (Fig 2B). To confirm that this decrease in WWP2 levels was not an artifact of HA-ubiquitin overexpression, we analyzed endogenous WWP2 levels in previously generated stable, polyclonal TMEM127 KO HEK293 cells (*5*) and observed an enrichment of WWP2 levels in TMEM127 KO-relative to the TMEM127 WT cells (Fig 3B). In addition, compared to single WWP2 KO cells in a background of endogenous TMEM127, double TMEM127 and WWP2 KO cells co-expressing a WWP2 construct showed significant decrease of the WWP2 construct levels (Fig 3C), in support of the post-transcriptional nature of the TMEM127-mediated WWP2 downregulation. Additionally, in agreement with the asymmetrical impact of TMEM127 and SUSD6 towards WWP2, we found that TMEM127 expression dominantly impacted WWP2 expression in contrast to SUSD6 (Fig 3D).

To better understand the requirements for the TMEM127 effect toward WWP2 autoubiquitination, we performed HA-ubiquitin co-immunoprecipitation assays with mutant versions of TMEM127. Based on our findings of HLA ubiquitination (Fig 2B), we hypothesized that TMEM127 Y236A would be deficient in eliciting WWP2 ubiquitin signals. Indeed, TMEM127 Y236, lacking PY-WW interactions, was incapable of promoting WWP2 ubiquitination to the same extent as WT TMEM127 (Fig 3E).

E3 ligase adaptors can also have roles in membrane protein trafficking of substrates and/or their cognate E3 ligases (*24*). We have previously shown that TMEM127 is expressed at multiple endomembrane domains and undergoes clathrin-mediated endocytosis, and that mutation targeting its endocytic domain leads to trapping of TMEM127 at the cell surface (*1, 3, 5*). We thus examined whether the TMEM127 endocytosis-deficient mutant L210A influenced WWP2 ubiquitination. The L210A mutant has an intact PY sequence so we first investigated its ability to interact with WWP2. We found that the L210A-WWP2 interaction is preserved and indistinguishable from WT TMEM127 (Fig 3F). Despite its ability to interact with WWP2, the TMEM127 L210A mutant was deficient in promoting WWP2 ubiquitination when compared with WT TMEM127 (Fig 3E), regardless of the addition of ectopic HLA-B to the complex (Fig 3G), suggesting that TMEM127 endocytic function is relevant for WWP2 autoubiquitination.

### TMEM127 recruits WWP2 to the cell surface through canonical PY and endocytic interactions

To investigate the contribution of protein localization to these findings, we next used confocal microscopy and analyzed the distribution of ectopic WWP2-Myc/Flag in double WWP2/TMEM127 KO HEK293 cells expressing GFP-only (TMEM127 KO) or GFP-tagged WT TMEM127. These experiments revealed that in cells expressing EV, or TMEM127 KO without GFP expression, WWP2 was diffusely distributed in the cytoplasm, while TMEM127 WT re-expression led to a marked re-distribution of WWP2 into punctate signals which strongly overlapped with the TMEM127 localization (Fig4 A). In addition, the overall WWP2 signals were markedly decreased in TMEM127 rescued cells (Fig 4A quantification plot). These results align with our observations of Fig 3B and 3C, suggesting that the effect of TMEM127 in reducing the WWP2 abundance is related to its intracellular redistribution. Next, we extended these analyses to the endocytic and PY-mutants. Neither expression of TMEM127 L210A nor TMEM127 Y236A were capable of rescuing the endomembrane-bound, punctate profile of WWP2 observed after WT TMEM127 rescue (Fig 4B), as reflected by decreased co-localization between WWP2 and each of these TMEM127 mutants (Fig 4B quantification plot). Additionally, we did not detect a measurable difference between TMEM127 L210A and Y236A in these assays, suggesting that both functional domains contribute to the WWP2 redistribution. These results support a model whereby TMEM127 expression leads to WWP2 protein downregulation, likely via its ubiquitination and cellular localization. This mechanism was dependent on TMEM127 endocytic competence and ability to physically interact with WWP2.

**Figure 4.**
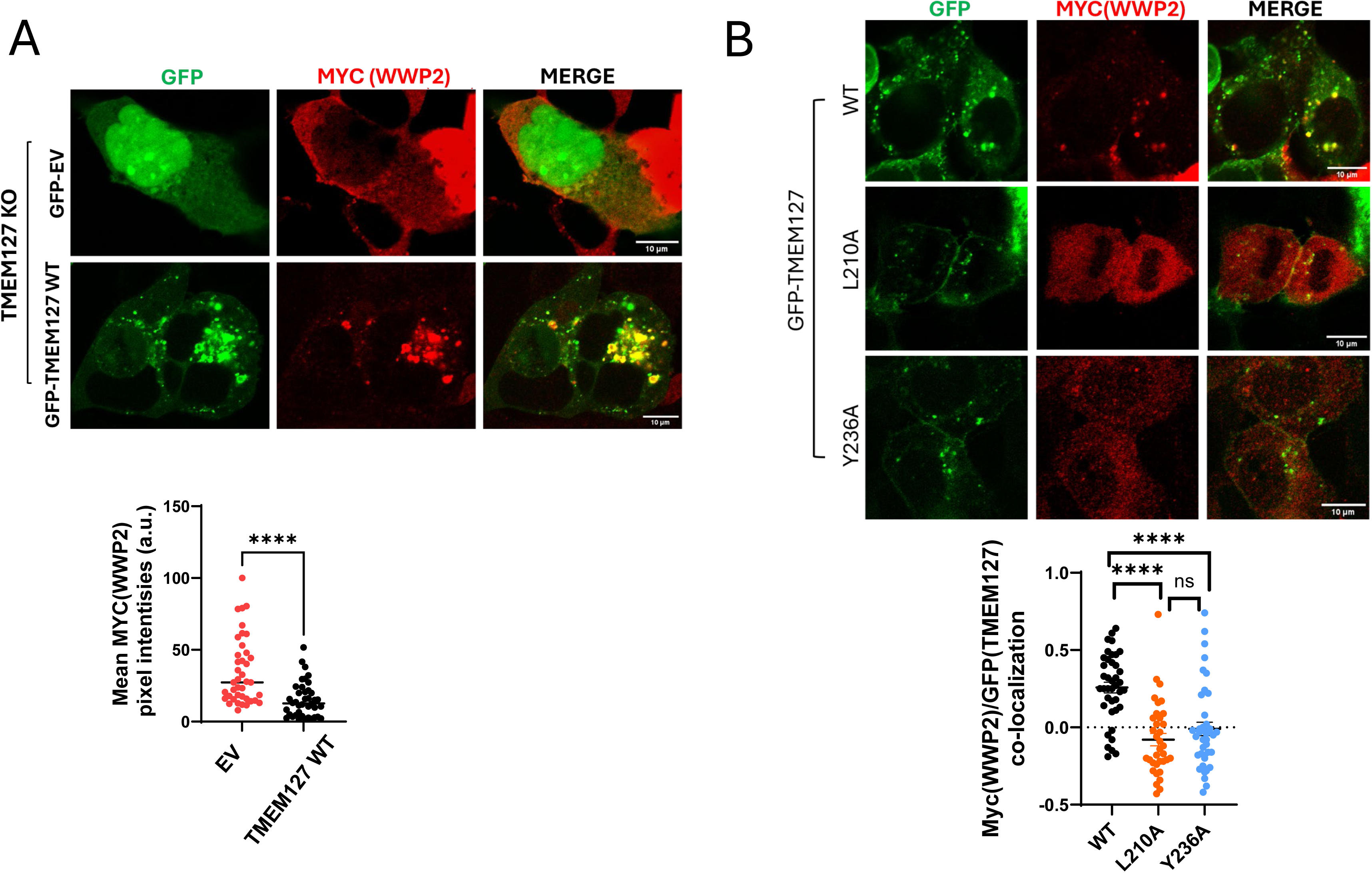
WWP2 subcellular distribution is impacted by TMEM127 expression. **A**) (Top) Representative images of confocal microscopy of WWP2 and TMEM127-depleted HEK293 cells co-expressing WWP2-Myc/Flag, and GFP-EV or GFP-TMEM127 WT constructs immunostained with a Myc (WWP2) antibody, pseudocolored in red. Scale bar is 10µm; (Bottom) Quantification of average WWP2 (Myc) signals in GFP+ cells from 3 independent experiments (n=80 cells; 40 cells/genotype), and compared using paired Student’s t test. N=3. **B)** (Top) Representative images of confocal microscopy of HEK293 cells co-expressing WWP2-Myc/Flag, and GFP-TMEM127 WT (N=40), L210A (N=35), and Y236A (N=40) constructs immunostained with a Myc (WWP2) antibody, pseudocolored in red. (Bottom) Quantification of colocalization analysis of Myc/GFP signals from captured images. A Pearson’s correlation value was calculated for each genotype and statistically analyzed using a paired Student’s t test. (****): p < 0.0001. Scale bar is indicated. N=3.

### WWP2 stabilizes the adaptor protein TMEM127 and SUSD6 in the ternary complex

The observations above suggested that, while TMEM127 and SUSD6 both contributed to the complex composition, their specific roles potentially differed. To further investigate the contribution of each adaptor to the complex assembly and stability, we performed co-IPs in cells expressing HLA-B and WWP2 in the presence of TMEM127 Y236A and found that, in addition to its inability to bind WWP2, TMEM127 Y236A also failed to bind SUSD6, while still maintaining interactions with HLA in this setting (Fig 5A). These findings were supported by reciprocal SUSD6 IP (Suppl. Fig 3A). TMEM127 Y236A was also capable of interacting with HLA alone (Fig 5B) and SUSD6 alone (Fig 5C). However, we found that SUSD6 interactions with TMEM127 were undetectable when HLA was co-expressed (Fig 5D), with reciprocal IPs confirming this observation (Suppl. Fig 3B). The TMEM127-SUSD6-HLA interactions were recovered when WWP2 was co-expressed (Fig 1C, 1E). Thus, we hypothesized that the TMEM127-SUSD6 interaction could be indirect, and dependent on WWP2. To test this hypothesis, we generated a SUSD6 mutant harboring a mutation on a PY-like motif (Y177A) that was recently reported to be critical for WWP2 interactions(*21*). After confirming that SUSD6 Y177A did not interact with WWP2 in our system (Suppl Fig 3C), we compared TMEM127-SUSD6 interactions in cells co-expressing HLA-B, WWP2, and SUSD6 WT or Y177A. Again, TMEM127-SUSD6 WT interactions were restored when WWP2 was added to the cells, but undetectable in cells expressing ectopic HLA-B only (Fig 5E). In contrast, TMEM127-SUSD6 Y177A interaction was not detectable, despite the presence of WWP2 (Fig 5E), suggesting that the TMEM127-SUSD6 interactions are unstable and dependent on WWP2. These findings show that the WWP2-SUSD6 interaction is required for the full STW assembly with its substrate MHCI.

**Figure 5.**
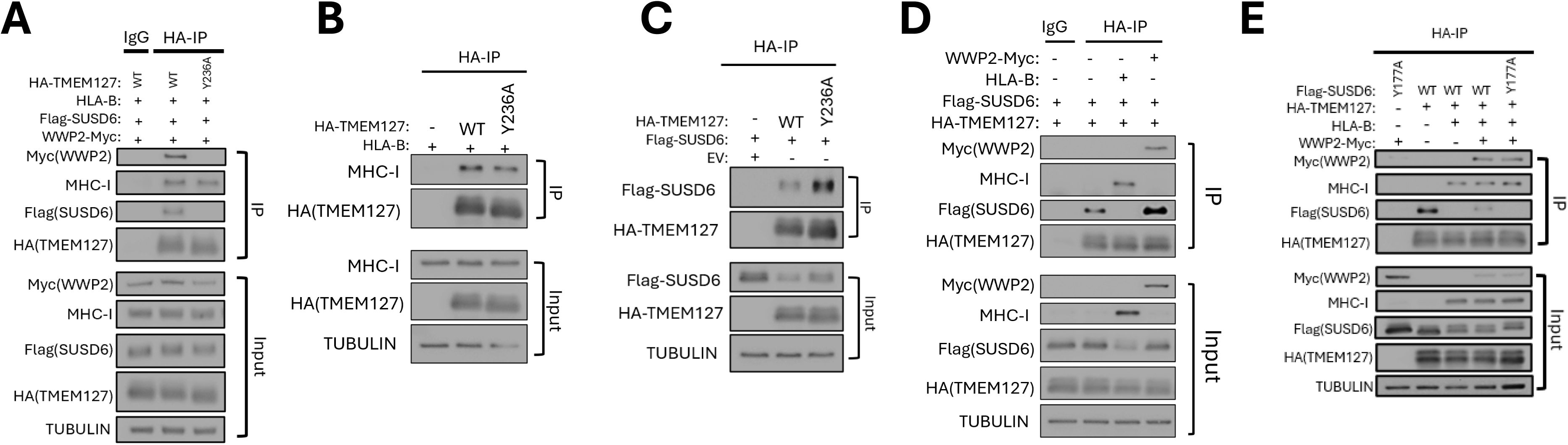
WWP2 stabilizes TMEM127 and SUSD6 in the ternary complex. **A)** Protein lysates from WWP2- and TMEM127-depleted HEK293 cells co-transfected with WWP2-Myc, HLA-B, Flag-SUSD6, and HA-TMEM127 were co-immunoprecipitated with a HA (TMEM127) or an IgG serotype control antibody and immunoprecipitates were immunoblotted for MHC-I, Flag (SUSD6), and Myc (WWP2). Corresponding whole cell lysate inputs were analyzed by immunoblotting with identical antibodies as co-IP blots; **B)** Protein lysates from WWP2- and TMEM127-depleted HEK293 cells co-transfected with HA-TMEM127 WT or HA-TMEM127 Y236A and HLA-B were co-immunoprecipitated with a HA (TMEM127) antibody and immunoprecipitates were immunoblotted for MHC-I. Corresponding whole cell lysate inputs were analyzed by immunoblotting with identical antibodies as co-IP blots; **C)** Protein lysates from WWP2- and TMEM127-depleted HEK293 cells co-transfected with HA-TMEM127 WT or HA-TMEM127 Y236A and Flag-SUSD6 were co-immunoprecipitated with a HA (TMEM127) antibody and immunoprecipitates were immunoblotted for Flag (SUSD6). Corresponding whole cell lysate inputs were analyzed by immunoblotting with identical antibodies as co-IP blots; **D)** Protein lysates from WWP2 and TMEM127-depleted HEK293 cells co-transfected with WWP2-Myc, HLA-B, Flag-SUSD6, and HA-TMEM127 WT were co-immunoprecipitated with an HA (TMEM127) or an IgG serotype control antibody and immunoprecipitates were immunoblotted for MHC-I, Flag (SUSD6), and Myc (WWP2). Corresponding whole cell lysate inputs were analyzed by immunoblotting with identical antibodies as co-IP blots. (Findings from these experiments are reproducible in different experiments within article); **E)** Protein lysates from WWP2 and TMEM127-depleted HEK293 cells co-transfected with WWP2-Myc, HLA-B, HA-TMEM127, and Flag-SUSD6 WT or Y177A mutant were co-immunoprecipitated with a HA (TMEM127) and immunoprecipitates were immunoblotted for MHC-I, Flag (SUSD6), and Myc (WWP2). Each experiment was performed 2-4 times.

Unexpectedly, we observed that the expression of Flag-SUSD6 was markedly reduced when HLA-B was overexpressed. A recent report suggested an inverse relationship between MHC-I and SUSD6 expression(*25, 26*). Therefore, we investigated whether this decline was due to association with WWP2, which might suggest that SUSD6 could also be a target of WWP2. However, we observed that HLA-B overexpression also downregulated SUSD6 Y177A, which is incapable of binding to WWP2 (Suppl Fig. 3D), suggesting that SUSD6 downregulation by HLA was independent of WWP2. Collectively, our data suggest that TMEM127 recruits WWP2-SUSD6 to the complex and promotes WWP2 and HLA ubiquitination. While the mechanism by which MHC-I expression influences SUSD6 level remains to be elucidated, we showed that it is unlikely to be mediated by WWP2.

### Modulation of MHC-I by TMEM127 is conserved across species and tissue types

Next, we investigated how TMEM127 loss impacted endogenous HLA expression. Treatment of TMEM127 WT and KO HEK293 cells with ribosomal inhibitor cycloheximide showed that HLA had an over 2-fold longer half-life in TMEM127 KO than in TMEM127 WT cells (Fig 6A). These results suggested that endogenous TMEM127 regulates endogenous MHC-I stability. We also found increased surface levels of endogenous MHC-I in TMEM127 KO cells by fluorescence activated cell sorting (FACS, Fig 6B), in agreement with earlier findings(*10*), an effect that was rescued in part by TMEM127 re-expression (Fig 6C).

**Figure 6.**
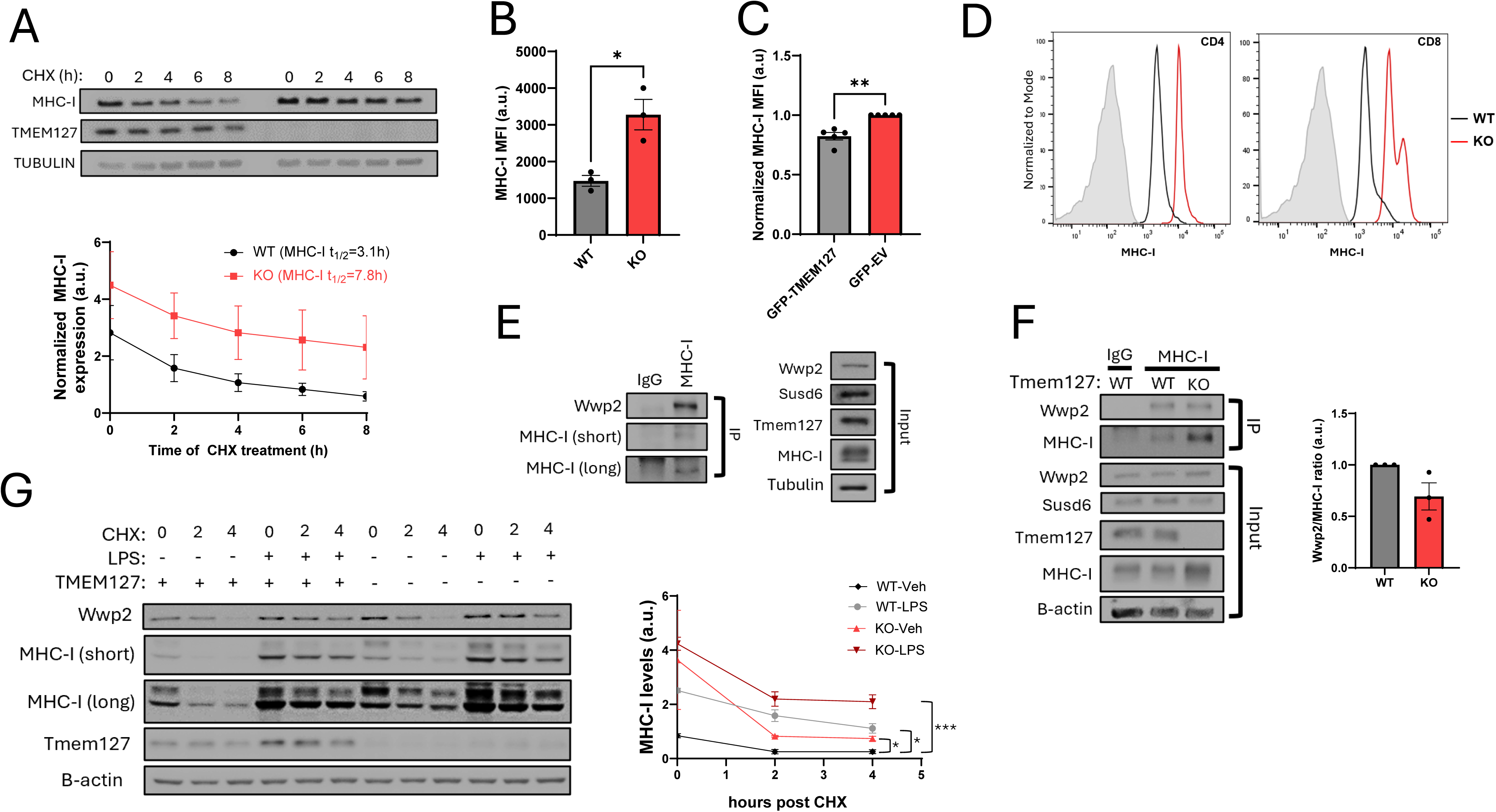
Modulation of MHC-I by TMEM127 is conserved across species and tissue types. **A)** Protein lysates from TMEM127 depleted HEK293 cells (and TMEM127 WT controls) treated with cycloheximide at 100ug/mL for 0, 2, 4, 6, and 8 hours were analyzed for MHC-I expression by immunoblotting with an MHC-I antibody. N=3, Plot depicts the quantification of MHC-I expression normalized by Tubulin loading control (** p<0.01, ANOVA). MHC-I decrease was statistically analyzed using a 2way ANOVA test with multiple comparisons. A phase decay curve was generated from the data.(**) = p < 0.005; **B)** TMEM127 WT and KO HEK293 cells were stained with an MHC-I antibody conjugated with PE and DAPI and were analyzed by FACS. Quantification of MHC-I median fluorescence intensity (MFI) is reported and was statistically analyzed using by an unpaired t test with Welch’s correction. (*): p < 0.05. N=3; **C)** TMEM127 KO HEK293 expressing a TMEM127 construct with a GFP-reporter or an GFP only empty vector control were stained with an MHC-I conjugated with PE and DAPI, and MHC-I (PE) signals were analyzed from GFP positive populations. MHC-I median fluorescence intensity (MFI) is reported as percent of ‘EV’ (TMEM127 KO) and was statistically analyzed by a paired two-tailed T test. (**): p < 0.005. N=5; **D)** Representative histograms of surface MHC-I expression in CD4+ and CD8+ T cells quantified by FACS in spleens of Tmem127 WT and KO mice (N=2/genotype); **E)** Protein lysates from the spleen of a Tmem127 WT mouse were co-immunoprecipitated with an MHC-I antibody or a serotype antibody control and immunoprecipitates were immunoblotted for WWP2. Corresponding whole cell lysate inputs were analyzed by immunoblotting with identical antibodies as co-IP blots. N=2; **F)** Protein lysates from the spleen of a Tmem127 WT or Tmem127 KO mouse were co-immunoprecipitated with an MHC-I antibody or a serotype antibody control and immunoprecipitates were immunoblotted for WWP2. Corresponding whole cell lysate inputs were analyzed by immunoblotting with identical antibodies as co-IP blots. N=6; 3 mice/Tmem127 genotype; **G)** Protein lysates from Tmem127 WT or KO RAW 264.7 cells stimulated with LPS at 100ng/mL for 24 hours and cycloheximide at 100ug/mL for 0, 2, and 4 hours were analyzed for Wwp2, MHC-I, and Tmem127 expression normalized by B-actin loading control. N=3; Quantification of MHC-I levels in WT and KO samples (Statistical analyses performed with 2-way ANOVA and Tukey’s multiple comparisons test (*p<0.05, *** p<0.001.

Germline loss-of-function mutations in *TMEM127* cause susceptibility to pheochromocytomas (PPGL), rare neuroendocrine adrenomedullary neoplasms(*1*). In *TMEM127* mutant PPGLs, the WT TMEM127 allele is deleted and the translated TMEM127 protein is unstable, quickly degraded, and thus not detectable by immunoblotting(*1, 5*) (Suppl Fig 4A). We investigated the expression of MHC-I in PPGLs and found that *TMEM127* mutant PPGLs showed a trend toward higher levels of HLA relative to other genotypes (Suppl. Fig 4A). Additionally, endogenous WWP2 expression, although detected at low levels in the PPGL tumor lysates, was highest in tumors harboring *TMEM127* mutations (Suppl. Fig 4A), mirroring the inverse relationship between TMEM127 and HLA or WWP2 expression (e.g. Fig 6A). These findings are consistent with the role of TMEM127 as a regulator of HLA and WWP2 levels across cell types. To expand on these findings, we next examined tissue from the whole-body *Tmem127* KO mice. These mice, which we reported earlier, are viable and have a normal lifespan(*27*). We found that MHC-I expression levels were higher in *Tmem127* KO relative to *Tmem127* WT across different tissues (Suppl. Fig 4B) and cell types (Fig 6D), in support of a species and tissue agnostic effect of TMEM127 towards MHC-I.

To provide physiological support for our ectopic co-IPs shown in Figs 1-3, we sought to detect the complex endogenously. Using lysates from the spleen of a *Tmem127* WT mouse, we immunoprecipitated MHC-I and found that endogenous Wwp2 was recovered from the immunoprecipitate, supporting that the interaction between Wwp2 and MHC-I exists physiologically (Fig 6E). We were also able to recover Wwp2 from endogenous MHC-I immunoprecipitates from lysates of *Tmem127* KO spleen, although at lower levels than those from WT spleen, when normalized by the abundance of the MHC-I pulldown (Fig 6F). As the Tmem127 KO mice have intact *Susd6* (Fig 6F), the decreased, but not absent, Wwp2 detected by MHC-I pulldown likely reflects the ability of Susd6 to bridge Wwp2-MHC-I interaction in *TMEM127*-deficient cells, similar to what we observed in the ectopic model (Fig 1C). Of note, in the KO spleen samples, MHC-I was higher both at the level of whole cell lysates and pulldown, which we attribute to lower MHC-I ubiquitination and degradation (Fig 6F). These results are consistent with previous observations in genetically modified cells(*10*).

Lastly, to examine potential signaling that could regulate MHC-I levels and influence the STW complex assembly, we took advantage of the physiological role of the Toll-like receptor 4 (TLR4) in engaging an inflammatory response that activates MHC-I in macrophages (*28–31*). To this end, we generated *Tmem127* KO murine immortalized macrophage RAW 264.7 cell lines, stimulated them with the TLR4 ligand lipopolysaccharide (LPS), and measured MHC-I levels in the presence and absence of cycloheximide to assess its stability. As expected, LPS exposure led to upregulation of MHC-I levels in control cells (Fig 6G). Tmem127 KO cells showed the expected increase of MHC-I at baseline, and significantly slower clearance of MHC-I levels (Fig 6G), recapitulating the findings in HEK293 cells (Fig 6A), and consistent with an effect of Tmem127 in MHC-I turnover (*10*). MHC-I accumulation in *Tmem127* deficient cells was further increased by LPS (Fig 6G, Suppl Fig 4C), suggesting that Tmem127 depletion and LPS induction synergize in augmenting MHC-I. Additionally, these experiments revealed that both Tmem127 mRNA and protein expression were significantly upregulated by LPS (Fig 6G, Suppl Fig 4D). Taking together with the impact of TMEM127 on MHC-I ubiquitination, these assays suggest the existence of a TMEM127-related negative feedback mechanism that limits inflammation. In this instance, MHC-I induction in association with TLR-4 engagement is restricted by increased expression of TMEM127, which promotes the ubiquitination and eventual degradation of MHC-I.

## Discussion

In this work, we describe the role by TMEM127 in the degradation complex targeting MHC-I (*10*). Our study supports a model whereby TMEM127-WWP2 interactions signal the complex activity. We found that, beyond bridging the proximity between WWP2 and HLA, interactions between TMEM127 and WWP2 result in the catalytic activation, as reported independently (*21*), and subsequent autoubiquitination of the WWP2, suggesting a potential feedback mechanism to limit complex activity. In exploring this mechanism further, we observed that the TMEM127-mediated WWP2 autoubiquitination was co-dependent on two distinct facets of TMEM127, namely its PY motif, the binding site to WWP2 (*21*), and its endocytic properties, a novel aspect of TMEM127 engagement with the complex. These findings point to the relevance of these two domains for functional TMEM127-WWP2 connections.

The negative impact of TMEM127 PY mutant towards WWP2 autoubiquitination is likely explained by the inability of this TMEM127 mutant to elicit PY-WW interaction-dependent allosteric changes that modify WWP2 from a ‘closed’ catalytically inactive conformation to an ‘open’ catalytically active state via its interactions with WWP2 WW domains, which is a classic mechanism of HECT E3 ligase regulation (*21–23, 32–35*). Our observations that an endocytic deficient TMEM127 mutant, which retains TMEM127 at the cell surface, also fails to elicit WWP2 autoubiquitination despite maintaining functional interactions with WWP2, supports a two-tiered mechanism of regulation by TMEM127 towards HECT E3 ligases that involves endocytosis. Indeed, HECT E3 ligases preferentially modify substrates with K63 ubiquitin linkages, which leads to endocytic trafficking and lysosomal degradation of the ubiquitinated substrate(*24, 36–38*).

How TMEM127 endocytosis contributes to WWP2 ubiquitination remains undetermined, but our observation that WWP2 subcellular distribution is disrupted by the TMEM127 endocytic deficient mutant may be related to this process. Endocytosis may involve mechanisms related to membrane curvature dynamics, as in the association between NEDD4L activation and the endocytosis regulatory FCHO2 at clathrin-coated pits (*39*). Loss of *TMEM127* was found to impair both plasma membrane dynamics and the assembly and maturation of clathrin-coated pits (*40*). Whether membrane curvature and/or clathrin-coated pit function are related to WWP2 interactions, and if TMEM127 endocytosis and WW-PY binding are coordinated to facilitate WWP2 structural positioning within the complex, its catalytic activation and/or its proximity to its substrate will require further work. Within the context of the STW complex, both functional PY and endocytic TMEM127 domains could lead to MHC-I ubiquitination by releasing WWP2 from its inhibitory conformation and leading to its catalytic activation. In turn, the augmented WWP2 autoubiquitination and its subsequent degradation could serve as a self-limiting mechanism to terminate its action toward the substrate, as in other HECT E3 ligases (*22, 41, 42*).

We propose that TMEM127-WWP2 functional interactions stabilize the assembly of the complex. Our data reveal that, in the absence of WWP2, the ternary TMEM127-SUSD6-HLA complex is undetectable. Based on these findings, we hypothesize that SUSD6 and HLA may compete for TMEM127 binding sites, with the interactions with HLA being stronger than those with SUSD6. The recruitment of WWP2 provides an alternate SUSD6-binding interface, which permits an indirect interaction between TMEM127 and SUSD6 in the presence of HLA and WWP2 to form the quaternary STW complex. Future biophysical and structural studies are warranted to directly resolve the intermolecular properties of the STW complex.

Our data also supports functional nuances in the roles of the two ‘adaptor’ components of STW, TMEM127 and SUSD6. We found that while either TMEM127 or SUSD6 were required for detectable WWP2-HLA interactions (*10*), we noted asymmetrical effects of these two membrane proteins toward releasing WWP2 activity and influencing its stability. In our model, TMEM127 alone, but not SUSD6 alone, was capable of eliciting WWP2 activation and autoubiquitination. The lack of SUSD6 effect on WWP2 ubiquitination agrees with a recent report (*21*). However, in that study, TMEM127 expression alone failed to promote WWP2 ubiquitination, unlike our findings. One possible reason for the discrepancy is that in those assays reported earlier the effect of TMEM127 was compared with a distinct and potent WWP2 adaptor protein, NFDIP2 (*21*). It is possible that differences in the degree of WWP2 disinhibition by these distinct adaptors may have obscured the detectability of TMEM127-driven signals.

We demonstrate that loss of *TMEM127* impacts MHC-I turnover, ubiquitination, and internalization in human and animal models, supporting a ubiquitous role of the STW complex across species and tissue. Our exploration of the impact of Tmem127 in an inflammatory model revealed additional facets of the complex. We found that TLR4-mediated MHC-I induction and stabilization, a mechanism that enhances peptide cross-presentation by MHC-I (*33*), parallels the impact of Tmem127 KO toward MHC-I abundance, although likely through distinct mechanisms, as LPS stimulation was able to further extend the stability of MHC-I in *Tmem127-*deficient macrophages. We also show that LPS promotes Tmem127 expression and stability, suggesting a negative feedback mechanism that limits TLR4-mediated MHC-I induction, whereby TMEM127 upregulation promotes the ubiquitination and eventual degradation of MHC-I. These findings also suggest that transcriptional factors downstream to TLR-4 (e.g., NF-KB, AP-1, IRF3) (*43*) may control TMEM127 expression, providing a potential regulatory signal for TMEM127 that should be further explored. Nevertheless, the observation that TLR4 signaling expands the impact of Tmem127 deficiency toward MHC-I expression suggests that distinct MHC-I enhancement strategies may be potentially combined to amplify MHC-I augmentation.

Of translational relevance, by demonstrating that the recruitment of WWP2 is necessary for TMEM127 and SUSD6 to remain in the complex, it is plausible that targeting WWP2 could augment MHC-I expression and antigen presentation. Pharmacological inhibition of HECT E3 ligases is feasible, and recently, WWP2 inhibition was reported to ameliorate chronic kidney injury in vitro and in vivo, supporting its potential use for improving antigen presentation (*44, 45*).

Our study has limitations. Future work should confirm interactions within the STW complex using additional endogenous models and structural studies. Similarly, as a putative cancer-relevant mechanism, the activity of the complex should be investigated in a tumor model. How precisely TMEM127 recognizes MHC-I and WWP2 remains to be determined. Our work does not explore MHC-I peptide loading nor specific antigen presentation, which are critical facets of MHC-I activity, thus, future work should evaluate the specific impact of TMEM127 deficiency in immunopeptidome changes (*14, 17*).

Collectively, these novel structural and regulatory properties of STW point to potential vulnerabilities that may be leveraged to augment MHC-I cell surface expression, improving antigen presentation and potentially anti-cancer immunity.

## Methods

### AlphaFold 3 modeling

Amino acid sequences of TMEM127, SUSD6, WWP2, HLA-B, and B2M were processed using AlphaFold3 multimer and visualized using PyMol.

### Generation of vector constructs

Existing TMEM127 and WWP2 DNA constructs were either previously reported(*1, 5, 8*) or engineered to harbor mutations of interest via PCR-based site-directed mutagenesis. Briefly, mutagenic primers were designed to introduce appropriate nucleotide changes to a template DNA construct, and these were incorporated into a PCR reaction using Phusion High-Fidelity DNA Polymerase (Thermo Scientific) where the extension time was adjusted to allow for complete amplification of the entire plasmid. Resulting amplicons were digested with DpnI (New England Biotechnology) restriction endonuclease to remove any trace of parental, methylated, template DNA. Next, purified PCR product was transformed to One-Shot Stbl3 (Thermo Scientific) competent cells, and resulting clones were screened for accurate mutagenesis by Sanger Sequencing (MCLAB, San Francisco, CA). DNA constructs harboring isolated WWP2 functional domains ‘C2’, ‘C2+WW1’, ‘C2+WW1-2’, ‘C2+WW1-3’, ‘C2+WW1-4’, and ‘HECT’ were generated by site-directed mutagenesis as above. Here, an amino acid downstream of a functional domain (or combination of functional domains) was mutagenized to a stop codon using a WWP2 construct with a dual C-terminus Myc/Flag tag as a template. For the ‘HECT’ construct, a subcloning strategy was employed the HECT coding sequence was amplified incorporating an AsiSI (New England Biolab) restriction site at the 5’ position. Next, the WWP2 coding sequence insert was released from the vector by digestion with AsiSI and NotI (New England Biolab), and the open vector was purified. Next, the HECT amplicon was digested with AsiSI and NotI, the digested product was used in a ligation reaction with the digested open vector. Competent cells were transformed with the ligation reaction, and transformants screened via restriction enzyme analysis and Sanger sequencing for presence of insert and correct sequence, respectively. The Flag-SUSD6 construct was a generous gift from Iannis Aifantis (*22*).

### CRISPR-Cas9 editing and retroviral/lentiviral transductions

CRISPR-Cas9 editing technology was used to genomically modify HEK-293FT and RAW 264.7 cell lines using human and mouse-specific TMEM127 guide RNAs as we reported(*1, 5, 8, 27*), or cloned into pLENTICRISPR V2 targeting WWP2 (*6*). Briefly, parental HEK293-FT cells were co-transfected with viral packaging constructs pMD2.G (Addgene 12259), psPAX2 (Addgene 12260) and the pLENTICRISPRv2 construct containing the specified guide RNA. Viral supernatants were harvested at 48h and 72h post-transfection and used to transduce target cells in the presence of polybrene at 8ug/mL (Thermo Scientific). 48-72h post-transduction, cells were selected either by treatment with puromycin or by FACS depending on the transduced construct selectable marker.

### Cell culture and transfections

HEK-293FT cell lines (Thermo Fisher/Invitrogen) were cultured and maintained in Dulbecco’s Modified Eagle’s Medium (DMEM) (Corning) supplemented with heat-inactivated 10% fetal bovine serum (Gibco) and 1% penicillin-streptomycin in a humidified incubator at 37C degrees and 5% CO_2_ partial pressure. Nucleic acid transfections were performed using Lipofectamine 2000 (Invitrogen) to form nucleic acid-lipid complexes which were added dropwise to HEK293FT cell lines in serum-free media. 4-6 hours post-transfection, transfection media was replaced with complete DMEM media.

### Immunoblotting

Whole cell lysates from cultured cells were prepared by first washing cells with ice-cold PBS to remove excess serum, followed by scraping cells in a non-denaturing lysis buffer composed of 1% NP40, 150mM NaCl, 50mM Tris-HCL, 10% glycerol, and 2mM EDTA. Cell lysates were incubated on ice for 30 minutes, and vortexed briefly every 5 minutes. Following incubation, lysates were cleared by centrifugation at 13,200 rpm for 20 minutes at 4C and supernatants were quantified for total protein concentration using a colorimetric protein assay (BioRad). Protein lysates were analyzed by sodium-dodecyl sulfate polyacrylamide gel electrophoresis (SDS-PAGE) followed by immunoblotting detection. Briefly, an aliquot of protein lysates was mixed with a beta-mercaptoethanol/SDS containing buffer, boiled for 5 minutes at 100C and loaded into gel. Gels were transferred to a polyvinylidene difluoride (PVDF) membrane and blocked for 60 minutes with a 5% non-fat milk solution in Tris-buffered saline with 0.1% Tween 20 (TBST). Following blocking, membranes were incubated overnight with primary antibodies at 4C. The following day, excess primary antibody was washed with TBST, and membranes were counterstained with a secondary antibody for 60 minutes at room temperature. Membranes were developed using an HRP-based colorimetric detection kit using an imager.

### Immunoprecipitation and co-Immunoprecipitation

Magnetic beads ‘A’ or ‘G’ (Dynabeads – Invitrogen) were washed twice with PBS with rotation for 15 minutes. Following washes, magnetic beads were incubated with 0.5-1ug of a primary antibody diluted in 1mL of PBS with 0.03% Tween 20 (PBST) for 4 hours at 4C with rotation. 0.5mg-1.0mg of protein lysate was pre-cleared with empty magnetic beads for 30 minutes. Pre-cleared lysates were then mixed with antibody-bound beads and incubated overnight at 4C with rotation. The next day, beads were washed six times with ice-cold protein lysis buffer (1% NP40, 150mM NaCL) with rotation for 15 minutes at room temperature. Following washes, beads were boiled with 50uL of 1X SDS buffer for 6 minutes at 100C, and IP lysates were analyzed by immunoblotting.

### RNA isolation and real-time PCR

Total RNA from RAW 264.7 cell lines was isolated using Qiagen reagent kit following the manufacturer’s protocol. RNA was converted into cDNA using the High-Capacity cDNA Reverse Transcription kit and random hexamers (ThermoFisher) according to the manufacturer’s instructions. Primer pairs were designed for mouse Tmem127 and beta-actin were as reported(*1*). SYBRGreen (Bio-Rad) was used for quantitative realtime PCR (qRT-PCR) in a QuantStudio6 instrument (ThermoFisher). Expression levels were calculated using the delta-delta Ct method.

### Fluorescence microscopy

For confocal microscopy analysis, HEK293 cells were transduced with lentiviral particles coding WWP2-Myc/Flag and transfected with GFP-TMEM127 WT, L210A, and Y236A constructs, with the addition of a GFP-only control. Next, cells were seeded onto coverslips previously coated with 0.15% gelatin for 15 minutes. Prior to fixation, coverslips were washed once with ice-cold PBS and were subsequently fixed with 4% paraformaldehyde (PFA) for 20 minutes. Following fixation, excess PFA was washed with PBS, and coverslips were permeabilized and blocked with a 0.1% Triton-X, 2% BSA, respectively, in PBS for 60 minutes. Following permeabilization and blocking, coverslips were stained with a Myc antibody (Santa Cruz mc-53584) at a 1:150 dilution (in 0.1% Triton-X, 2% BSA buffer) overnight at 4C. The next day, coverslips were washed in PBS three times for 5 minutes, and counter-stained with Alexa Fluor 647 (Thermo Fisher Scientific) for 60 minutes at room temperature. Following counter-staining, coverslips were washed three times for 5 minutes with PBS at room temperature. Next, coverslips were stained with DAPI (Thermo Fisher Scientific) at a 1:1000 dilution (in PBS) for 5 minutes at room temperature. Excess DAPI was washed once with PBS, and coverslips were then mounted on glass slides with antifade mounting media (Vector Labs). Coverslips were imaged using a Zeiss LSM980 with an Airyscan 2 super-resolution instrument using a 63X oil-immersion objective. Colocalization analysis of WWP2 (Myc) and GFP (TMEM127) signals was performed using FIJI (fiji.sc) using the CoLoc2 plugin to generate a Pearson’s correlation coefficient from overlapping signals of cells positive for both Myc and GFP signals. Results were statistically analyzed using an unpaired Student’s t-test in the GraphPad Prism software (version 10.6.1.).

### Flow cytometry

For FACS analyses, HEK293 cells were trypsinized and washed once with ice-cold PBS, centrifuged, and resuspended in ice cold PBS + 4%FBS. Next, cells were stained for 45’ on ice with a PE-conjugated HLA A/B/C antibody (Clone W6/32, Biolegend, #311406). Viable Tmem127 KO or WT control spleen cells were resuspended in 100 μl of FACS buffer with 1 μl of Fc Block (anti-mouse Fc block CD16/32, catalog no. 101302, Biolegend) for 20 min at 4°C. Cells were then washed with FACS buffer and stained with surface marker antibodies targeting MHC-I (Biolegend cat#114614) and, to detect the respective T cell populations, CD4 (Biologend cat#116023) and CD8 (Biolegend cat#100712). Following the immunostaining, cells were washed extensively with PBS + 2%FBS. Live/dead staining was done with DAPI and cells were analyzed using a BD LSR-II instrument. Data was analyzed and displayed using FlowJo under an institutional license.

## Acknowledgements

We are grateful to Ianis Aifantis for providing the FLAG-SUSD6 construct and also to current and former members of the Dahia lab for their input and technical assistance. This work was supported by funds from the NIH National Cancer Institute (NCA) CA264468-S1 (H.G.C) CA264468-S2 (V.N.C.), CA264248, Paradifference Foundation, UT System Star Awards and Mike Hogg Fund (P.L.M.D). Confocal images were generated in the Core Optical Imaging Facility which is supported by UT Health San Antonio and NIH-NCI P30 CA54174. The Zeiss LSM confocal with Airyscan 2 was funded by NIH S10 grant 1S10OD036297-01. Flow cytometry analyses were undertaken at the Flow Cytometry Shared Resource at UT Health San Antonio which is supported by a grant from the National Cancer Institute (P30CA054174) to the Mays Cancer Center, a grant from the Cancer Prevention and Research Institute of Texas (CPRIT) (RP210126), a grant from the National Institutes of Health (S10OD030432), and support from the Office of the Vice President for Research at UT Health San Antonio. The studies were also supported by the Mays Cancer Center Drug Discovery Shared Resource (including the Target Discovery Core and the Center for Innovative Drug Discovery), which is funded by grants from NCI (P30CA054174) and CPRIT grants (RP210208 and RP250601).

## Author Contributions

H.G.C. designed, executed, analyzed experiments, and wrote the manuscript. V.N.C., S.Y.M., K.J., C.J., N.R., A.M., Y.X. performed experiments. Y.X., D.Z., R.C.T.A., P.L.M.D. discussed and analyzed the data. P.L.M.D. conceived and oversaw the study, co-wrote the manuscript and secured funding support. All authors reviewed and approved the manuscript.

---

**Supplemental Figure 1 (Related to Fig 1).**
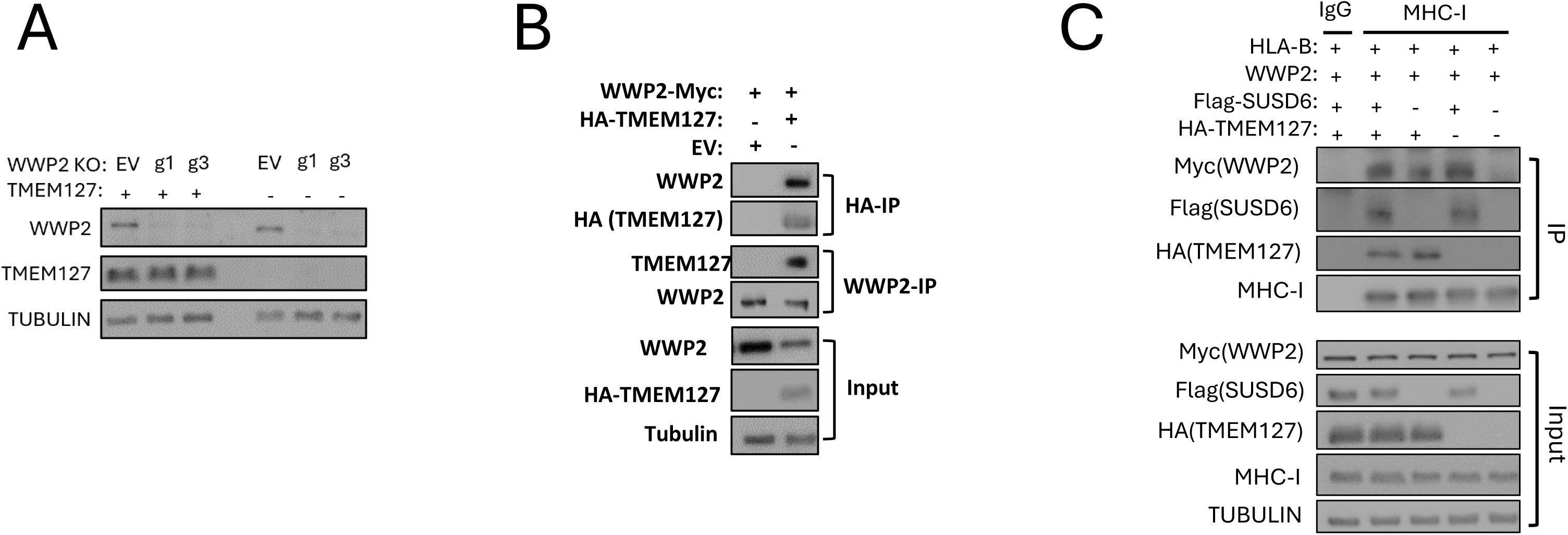
**A)** Protein lysates from WWP2 KO (‘g1’ and ‘g3’ refer to separate CRISPR guide RNAs used) TMEM127 WT and WWP2 KO, TMEM127 KO HEK293 cells were immunoblotted for WWP2, TMEM127, and TUBULIN, as a loading control; **B)** Protein lysates from WWP2 and TMEM127-depleted HEK293 cells co-transfected with WWP2-Myc and HA-TMEM127 were co-immunoprecipitated with HA (top) or WWP2 (middle) antibody and immunoprecipitates were immunoblotted for WWP2 and HA (TMEM127). Corresponding whole cell lysate inputs were analyzed by immunoblotting with identical antibodies as co-IP blots; **C)** Protein lysates from WWP2 and TMEM127-depleted HEK293 cells co-transfected with WWP2-Myc/Flag, HLA-B, Flag-SUSD6, and HA-TMEM127 were co-immunoprecipitated with an MHC-I or an IgG serotype control antibody and immunoprecipitates were immunoblotted for Myc (WWP2), Flag (SUSD6), HA (TMEM127). Corresponding whole cell lysate inputs were analyzed by immunoblotting with identical antibodies as co-IP blots.

**Supplemental Figure 2 (Related to Fig 3).**
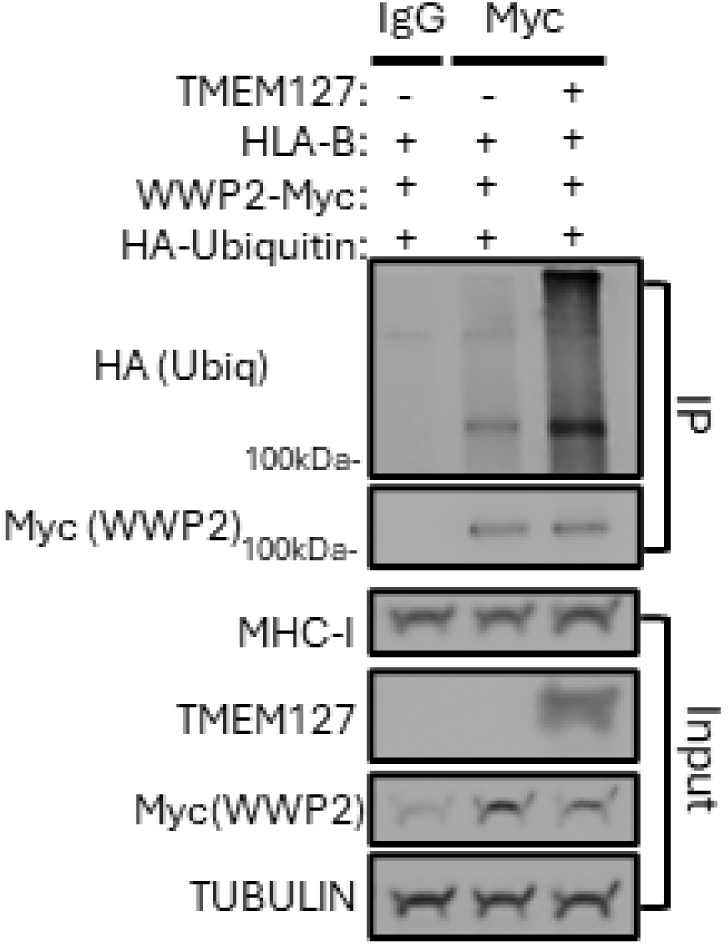
Protein lysates from WWP2 and TMEM127-depleted HEK293 cells co-transfected with WWP2-Myc, HLA-B, HA-ubiquitin, and TMEM127 were co-immunoprecipitated with a Myc (WWP2) or an IgG serotype control antibody and immunoprecipitates were immunoblotted for Myc (WWP2), and HA (ubiquitin). Corresponding whole cell lysate inputs were analyzed by immunoblotting with identical antibodies as co-IP blots. N=3

**Supplemental Figure 3 (Related to Fig 5).**
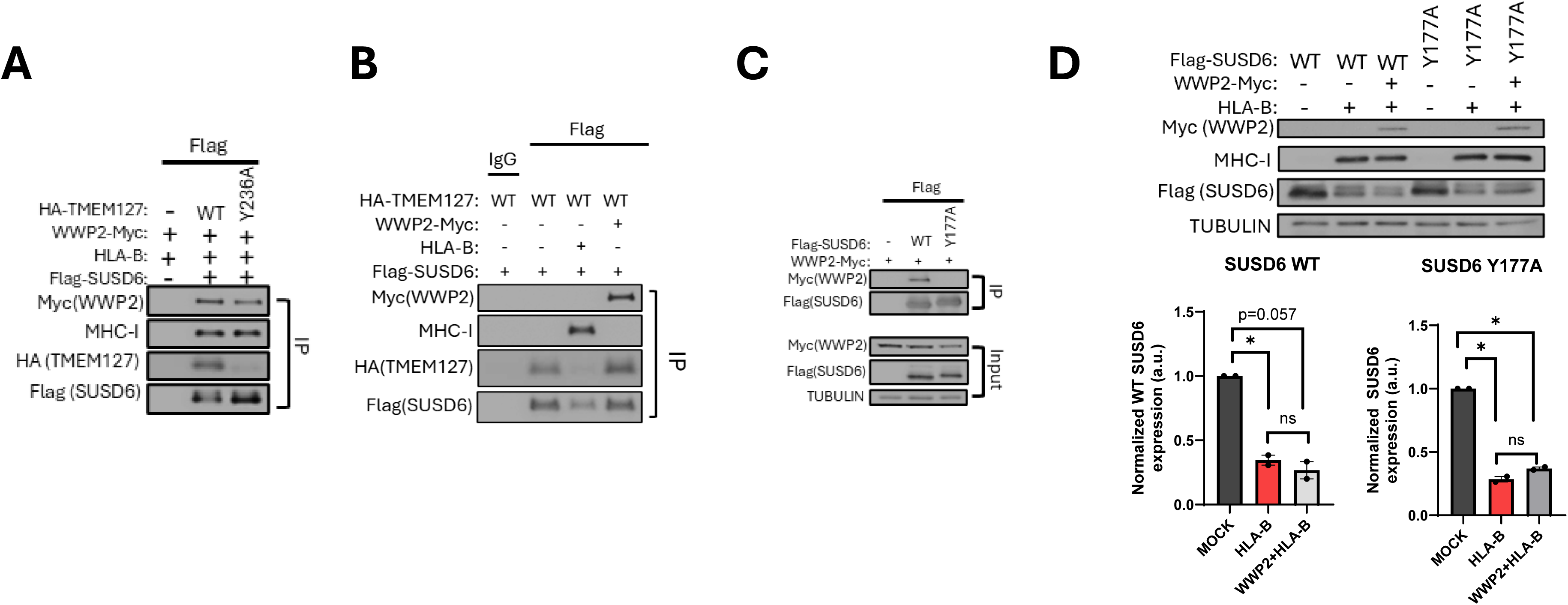
**A)** Protein lysates from WWP2 and TMEM127-depleted HEK293 cells co-transfected with WWP2-Myc, HLA-B, Flag-SUSD6, and HA-TMEM127 WT or Y236A mutant were co-immunoprecipitated with a Flag (SUSD6) and immunoprecipitates were immunoblotted for MHC-I, Flag (SUSD6), and HA (TMEM127). Corresponding whole cell lysate inputs were analyzed by immunoblotting with identical antibodies as co-IP blots; **B)** Protein lysates from WWP2 and TMEM127-depleted HEK293 cells co-transfected with WWP2-Myc, HLA-B, Flag-SUSD6, and HA-TMEM127 WT were co-immunoprecipitated with a Flag (SUSD6) or an IgG serotype control antibody and immunoprecipitates were immunoblotted for MHC-I, HA (TMEM127), and Myc (WWP2). Corresponding whole cell lysate inputs were analyzed by immunoblotting with identical antibodies as co-IP blots; **C)** Protein lysates from WWP2 and TMEM127-depleted HEK293 cells co-transfected with WWP2-Myc, HLA-B, Flag-SUSD6 WT or SUSD6 Y177A co-immunoprecipitated with a Flag (SUSD6) and immunoblotted for Flag (SUSD6), and Myc (WWP2). Corresponding whole cell lysate inputs were analyzed by immunoblotting with identical antibodies as co-IP blots. **D)** (Left) Protein lysates from WWP2 and TMEM127-depleted HEK293 cells co-transfected with WWP2-Myc, HLA-B, and Flag-SUSD6 WT or Y177A mutant were analyzed for Flag-SUSD6 expression by immunoblotting with a Flag (SUSD6) antibody. (Right) Quantification of Flag-SUSD6 bands normalized by Tubulin. Data is represented as percent of ‘MOCK’ and was statistically analyzed using an unpaired t test with Welch’s correction. (*): p<0.05. N=2.

**Supplemental Figure 4 (Related to Fig 6).**
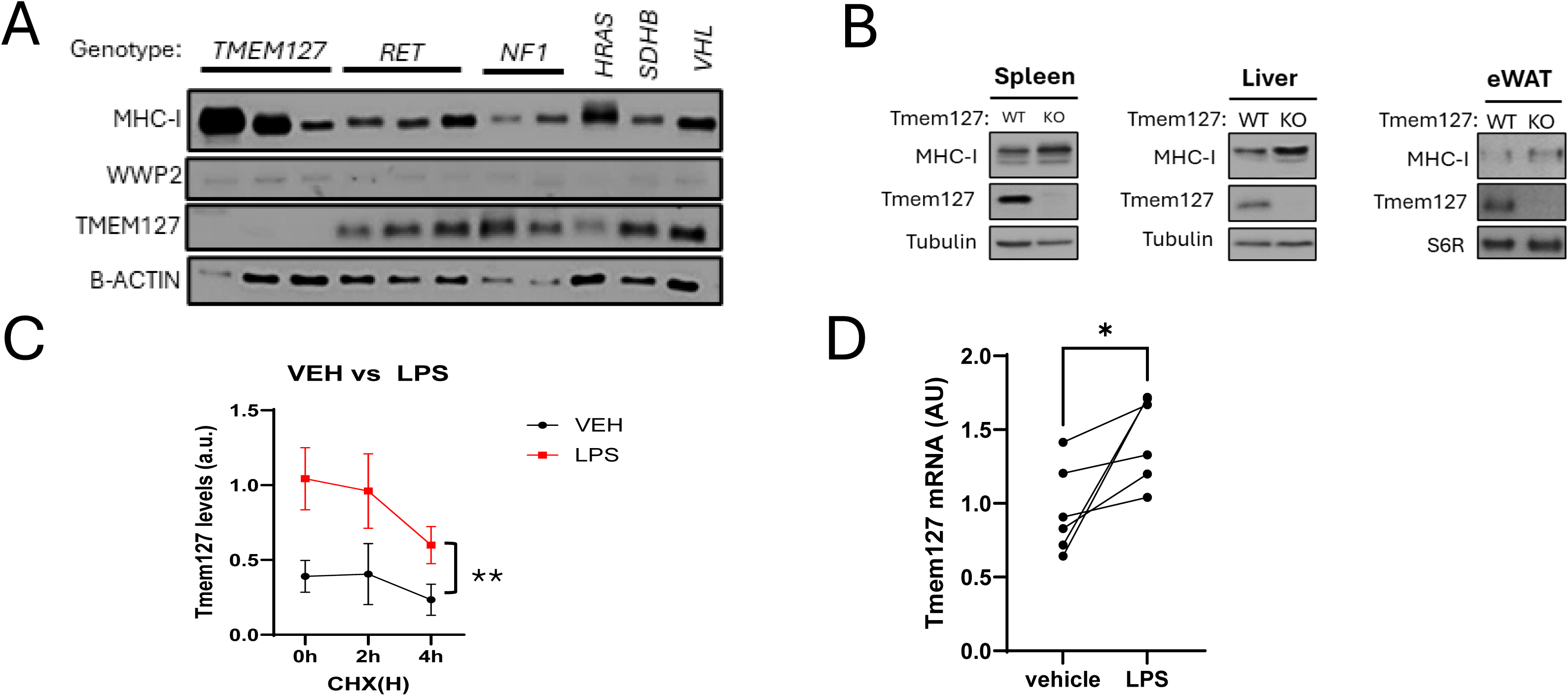
**A)** Protein lysates from pheochromocytomas and paragangliomas (PPGLs) of the indicated genotypes: *TMEM127, RET,* and *NF1* driving mutations (*RET, NF1, HRAS, SDHB and VHL-*mutant tumors express intact WT TMEM127) were analyzed for WWP2 and MHC-I expression by immunoblotting; **B)** Protein lysates from spleen, liver, and epididymal white adipose tissue (eWAT) from Tmem127 WT and KO of different genders and ages were analyzed for MHC-I expression by immunoblotting; Statistical analyses were performed by timepoint by paired (same genotype) or unpaired (distinct genotypes) Student’s t test. (*): p < 0.05, (**): p < 0.005, n.s.=nonsignificant. N=3; **C)** Quantification of Tmem127 protein levels from Tmem127 WT vehicle vs LPS treated groups. Tmem127 bands were normalized by B-actin loading control; Statistical analyses were performed by ANOVA, **p<0.005. N=3; D) Tmem127 mRNA expression after control (vehicle) or LPS 100ng/mL for 24 hours, quantified by real time PCR and normalized by b-actin (data are expressed as the mean of triplicate samples from 6 biological replicates), *= p<0.01, two-tailed paired Student’s t-test.

## References

1. Y. Qin et al., Germline mutations in TMEM127 confer susceptibility to pheochromocytoma. Nat Genet 42, 229–233 (2010).

2. L. Yao et al., Spectrum and prevalence of FP/TMEM127 gene mutations in pheochromocytomas and paragangliomas. JAMA 304, 2611–2619 (2010).

3. Y. Qin et al., The tumor susceptibility gene TMEM127 is mutated in renal cell carcinomas and modulates endolysosomal function. Hum Mol Genet 23, 2428–2439 (2014).

4. Y. Deng et al., Molecular and phenotypic evaluation of a novel germline TMEM127 mutation with an uncommon clinical presentation. Endocr Relat Cancer 25, X3 (2018).

5. S. K. Flores et al., Functional Characterization of TMEM127 Variants Reveals Novel Insights into Its Membrane Topology and Trafficking. J Clin Endocrinol Metab 105, e3142–3156 (2020).

6. E. Alix et al., The Tumour Suppressor TMEM127 Is a Nedd4-Family E3 Ligase Adaptor Required by Salmonella SteD to Ubiquitinate and Degrade MHC Class II Molecules. Cell Host Microbe 28, 54–68 e57 (2020).

7. P. L. Dahia, Pheochromocytoma and paraganglioma pathogenesis: learning from genetic heterogeneity. Nature reviews. Cancer 14, 108–119 (2014).

8. Q. Guo et al., TMEM127 suppresses tumor development by promoting RET ubiquitination, positioning, and degradation. Cell Rep 42, 113070 (2023).

9. D. Dersh et al., Genome-wide Screens Identify Lineage- and Tumor-Specific Genes Modulating MHC-I- and MHC-II-Restricted Immunosurveillance of Human Lymphomas. Immunity 54, 116–131 e110 (2021).

10. X. Chen et al., A membrane-associated MHC-I inhibitory axis for cancer immune evasion. Cell 186, 3903–3920 e3921 (2023).

11. S. Jhunjhunwala, C. Hammer, L. Delamarre, Antigen presentation in cancer: insights into tumour immunogenicity and immune evasion. Nat Rev Cancer 21, 298–312 (2021).

12. N. Xie et al., Neoantigens: promising targets for cancer therapy. Signal Transduct Target Ther 8, 9 (2023).

13. K. L. Rock, E. Reits, J. Neefjes, Present Yourself! By MHC Class I and MHC Class II Molecules. Trends Immunol 37, 724–737 (2016).

14. E. Shklovskaya, H. Rizos, MHC Class I Deficiency in Solid Tumors and Therapeutic Strategies to Overcome It. Int J Mol Sci 22, (2021).

15. A. M. Cornel, I. L. Mimpen, S. Nierkens, MHC Class I Downregulation in Cancer: Underlying Mechanisms and Potential Targets for Cancer Immunotherapy. Cancers (Basel) 12, (2020).

16. C. Galassi, T. A. Chan, I. Vitale, L. Galluzzi, The hallmarks of cancer immune evasion. Cancer Cell 42, 1825–1863 (2024).

17. D. Hwang et al., HLA-Shuttle: A system for enhancing antigen presentation in immunologically cold tumors. Sci Adv 12, eaeb0821 (2026).

18. J. Wang, Q. Lu, X. Chen, I. Aifantis, Targeting MHC-I inhibitory pathways for cancer immunotherapy. Trends Immunol 45, 177–187 (2024).

19. J. Abramson et al., Accurate structure prediction of biomolecular interactions with AlphaFold 3. Nature 630, 493–500 (2024).

20. D. Rotin, S. Kumar, Physiological functions of the HECT family of ubiquitin ligases. Nat Rev Mol Cell Biol 10, 398–409 (2009).

21. S. V. Blundell et al., Mammalian and bacterial adaptors function as co-disinhibitory pairs to activate the E3 ubiquitin ligase WWP2. J Biol Chem 301, 110847 (2025).

22. Z. Chen et al., A Tunable Brake for HECT Ubiquitin Ligases. Mol Cell 66, 345–357 e346 (2017).

23. H. Jiang, S. N. Thomas, Z. Chen, C. Y. Chiang, P. A. Cole, Comparative analysis of the catalytic regulation of NEDD4-1 and WWP2 ubiquitin ligases. J Biol Chem 294, 17421–17436 (2019).

24. S. Leon, R. Haguenauer-Tsapis, Ubiquitin ligase adaptors: regulators of ubiquitylation and endocytosis of plasma membrane proteins. Exp Cell Res 315, 1574–1583 (2009).

25. H. Estephan et al., Hypoxia promotes tumor immune evasion by suppressing MHC-I expression and antigen presentation. EMBO J 44, 903–922 (2025).

26. R. Yang et al., Melanoma MHC-I-membrane-encapsulated Cu@ferrihydrite induces ferroptosis/cuproptosis and systematic immunity against tumor. J Control Release 388, 114281 (2025).

27. S. Srikantan et al., The tumor suppressor TMEM127 regulates insulin sensitivity in a tissue-specific manner. Nat Commun 10, 4720 (2019).

28. W. Yan et al., Lipopolysaccharide (LPS)-induced inflammation in RAW264.7 cells is inhibited by microRNA-494-3p via targeting lipoprotein-associated phospholipase A2. Eur J Trauma Emerg Surg 50, 3289–3298 (2024).

29. A. Ciesielska, M. Matyjek, K. Kwiatkowska, TLR4 and CD14 trafficking and its influence on LPS-induced pro-inflammatory signaling. Cell Mol Life Sci 78, 1233–1261 (2021).

30. P. Nair-Gupta et al., TLR signals induce phagosomal MHC-I delivery from the endosomal recycling compartment to allow cross-presentation. Cell 158, 506–521 (2014).

31. J. Mathé, M. Benhammadi, K. S. Kobayashi, S. Brochu, C. Perreault, Regulation of MHC Class I Expression in Lung Epithelial Cells during Inflammation. J Immunol 208, 1021–1033 (2022).

32. S. S. Shah, S. Kumar, Adaptors as the regulators of HECT ubiquitin ligases. Cell Death Differ 28, 455–472 (2021).

33. Z. Wang et al., A multi-lock inhibitory mechanism for fine-tuning enzyme activities of the HECT family E3 ligases. Nat Commun 10, 3162 (2019).

34. T. Mund et al., Disinhibition of the HECT E3 ubiquitin ligase WWP2 by polymerized Dishevelled. Open Biol 5, 150185 (2015).

35. T. Mund, H. R. Pelham, Control of the activity of WW-HECT domain E3 ubiquitin ligases by NDFIP proteins. EMBO Rep 10, 501–507 (2009).

36. J. Weber, S. Polo, E. Maspero, HECT E3 Ligases: A Tale With Multiple Facets. Front Physiol 10, 370 (2019).

37. H. C. Kim, J. M. Huibregtse, Polyubiquitination by HECT E3s and the determinants of chain type specificity. Mol Cell Biol 29, 3307–3318 (2009).

38. J. A. Nathan, H. T. Kim, L. Ting, S. P. Gygi, A. L. Goldberg, Why do cellular proteins linked to K63-polyubiquitin chains not associate with proteasomes? EMBO J 32, 552–565 (2013).

39. Y. Sakamoto et al., The Nedd4L ubiquitin ligase is activated by FCHO2-generated membrane curvature. EMBO J 43, 5883–5909 (2024).

40. T. J. Walker et al., Loss of tumor suppressor TMEM127 drives RET-mediated transformation through disrupted membrane dynamics. Elife 12, (2024).

41. D. Sinha, S. S. Shah, S. Kumar, HECT ubiquitin ligases as regulators of inflammatory signalling. Cell Death Differ, (2026).

42. T. Perron et al., CYYR1 promotes the degradation of the E3 ubiquitin ligase WWP1 and is associated with favorable prognosis in breast cancer. J Biol Chem 300, 107601 (2024).

43. E. M. Palsson-McDermott, L. A. O’Neill, Signal transduction by the lipopolysaccharide receptor, Toll-like receptor-4. Immunology 113, 153–162 (2004).

44. T. Mund, M. J. Lewis, S. Maslen, H. R. Pelham, Peptide and small molecule inhibitors of HECT-type ubiquitin ligases. Proc Natl Acad Sci U S A 111, 16736–16741 (2014).

45. M. Wu et al., Targeting renal tubular WWP2 to restore mitochondrial OXPHOS integrity retards the AKI-to-CKD transition. Mol Ther 34, 1576–1596 (2026).

